# From features to slice: parameter-cloud modeling of spatial transcriptomics for simulation and 3D interpolatory augmentation

**DOI:** 10.64898/2025.12.04.692251

**Authors:** Yiru Chen, Manfei Xie, Yunfei Hu, Weiman Yuan, Hirak Sarkar, Bingshan Li, Lu Zhang, Xin Maizie Zhou

**Affiliations:** Systems and Informatics of Zhejiang University-University of Edinburgh Institute, Zhejiang University School of Medicine, Hangzhou, Zhejiang, China; Department of Biomedical Engineering, Vanderbilt University, USA; Department of Computer Science, Vanderbilt University, USA; College of Connected Computing, Vanderbilt University, USA; Department of Molecular Physiology and Biophysics, Vanderbilt University, USA; Department of Computer Science, Hong Kong Baptist University, Hong Kong

**Keywords:** Spatial Transcriptomics, ST Simulation, Computational Interpolation, 3D Reconstruction

## Abstract

Computational simulation and data augmentation of spatial transcriptomics (ST) are essential for quantitative benchmarking, reproducibility, and methodological innovation. Yet current models often lack flexibility in controlling spatial and transcriptional heterogeneity, fail to capture higher-order gene dependencies, and rarely extend to three-dimensional or alignment-aware contexts. Here we present FEAST, a computational infrastructure that models ST data within a parameter cloud - a latent manifold encoding gene-level mean, variance, and sparsity. By sampling and perturbing this manifold, FEAST generates high-fidelity synthetic slices with tunable spatial and transcriptional variation, enabling systematic evaluation of clustering, deconvolution, and spatial alignment algorithms. Beyond two dimensions, FEAST performs 3D parameter-cloud interpolation guided by optimal transport and benchmark alignment, reconstructing continuous tissue architectures while preserving molecular coherence. Together, these capabilities establish FEAST as a foundational platform for standardized benchmarking, data augmentation, and 3D reconstruction in spatial transcriptomics. The source code and tutorials for FEAST are publicly available at https://github.com/maiziezhoulab/FEAST, and can be installed via Pypi at https://pypi.org/project/FEAST-py/.

## 1 Introduction

Spatial transcriptomics (ST) technologies have revolutionized transcriptomic profiling by capturing gene expression within its native tissue context.^1–7^ By combining molecular readouts with spatial localization, ST enables gene expression maps that reveal the organization of transcriptional programs across tissue architecture, providing new insights into biological topics like development, neural circuitry, and disease. Alongside these experimental breakthroughs, a rapidly expanding ecosystem of computational methods has emerged to analyze ST data, encompassing spatial domain identification,^8–11^ cell-type deconvolution,^12^ multislice alignment and integration,^13–15^ and three-dimensional (3D) tissue reconstruction.^16, 17^

As these analytical frameworks proliferate, the lack of standardized, controllable, and interpretable simulation benchmarks has become a critical bottleneck for rigorous validation and reproducibility.^18^ In silico simulation provides a unique avenue for generating spatially resolved ground-truth datasets with fully known parameters, enabling systematic evaluation of algorithmic performance, robustness, and scalability.^19, 20^ Existing ST simulators remain limited in scope. Reference-based models such as SRTsim preserve spatial expression patterns from specific datasets but fail to generalize across tissues with distinct geometries or cellular compositions.^21–24^ Multimodal simulators like scDesign3 capture inter-modality correlations but are computationally intensive and difficult to scale to large modern ST datasets.^25^ Deep-learning simulators such as scCube leverage extensive training data yet lack interpretability and fine control over biological perturbations.^22^ De novo approaches such as scMultiSim can generate synthetic tissues from regulatory priors but remain challenging to calibrate for routine benchmarking.^26^

Additionally, a further gap lies in dimensionality: most simulators generate only two-dimensional (2D) slices, whereas current research increasingly demands quantitative evaluation of 3D reconstruction and spatiotemporal alignment methods.^16, 27^ Without realistic volumetric ground truths, assessment of 3D alignment and interpolation algorithms remains ad hoc and dataset-specific. This is more like a data augmentation problem: so how to use the most of these limited slices, to reconstruct realistic and biologically meaningful 3D region becomes to be a critical computational issue.

To overcome these challenges, we introduce FEAST (FEAture-space based modeling for Spatial Transcriptomics), a computational framework that models spatial transcriptomics data within a parameter cloud - a latent manifold capturing gene-level mean, variance, and sparsity. By sampling and perturbing this manifold, FEAST generates high-fidelity synthetic slices with tunable transcriptional and spatial alterations, enabling quantitative benchmarking of diverse analytical tasks including clustering, deconvolution, and spatial alignment. Beyond 2D simulation, FEAST performs 3D parameter-cloud interpolation across adjacent slices, integrating optimal transport and alignment-guided coordinate mapping to reconstruct missing sections and continuous 3D architectures for various downstream analyses. This process also supports controlled geometric perturbation and benchmark alignment, allowing quantitative evaluation of registration and reconstruction algorithms under known transformations.

Currently, FEAST is focusing on establishing a flexible and biologically interpretable simulation for comprehensive benchmarking and augmentation of spatial transcriptomics analyses. We anticipate that FEAST will serve as a reference framework for standardized 3D benchmarking simulation and data augmentation, providing a foundation for the next generation of computational spatial transcriptomics, and may be an extendable model for other tasks.

## 2 Methods

### 2.1 Single-slice simulation

To generate realistic ST data that preserve the distributional characteristics observed in real slices, FEAST builds a hierarchical generative framework operating at the gene level (Fig. 1). Specifically, each gene *g* is represented by a parameter vector 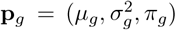, capturing its mean, variance, and sparsity (zero proportion) within a single ST slice. These parameters are learned from empirical data and then used to simulate new, biologically plausible expression profiles.

**Fig. 1.**
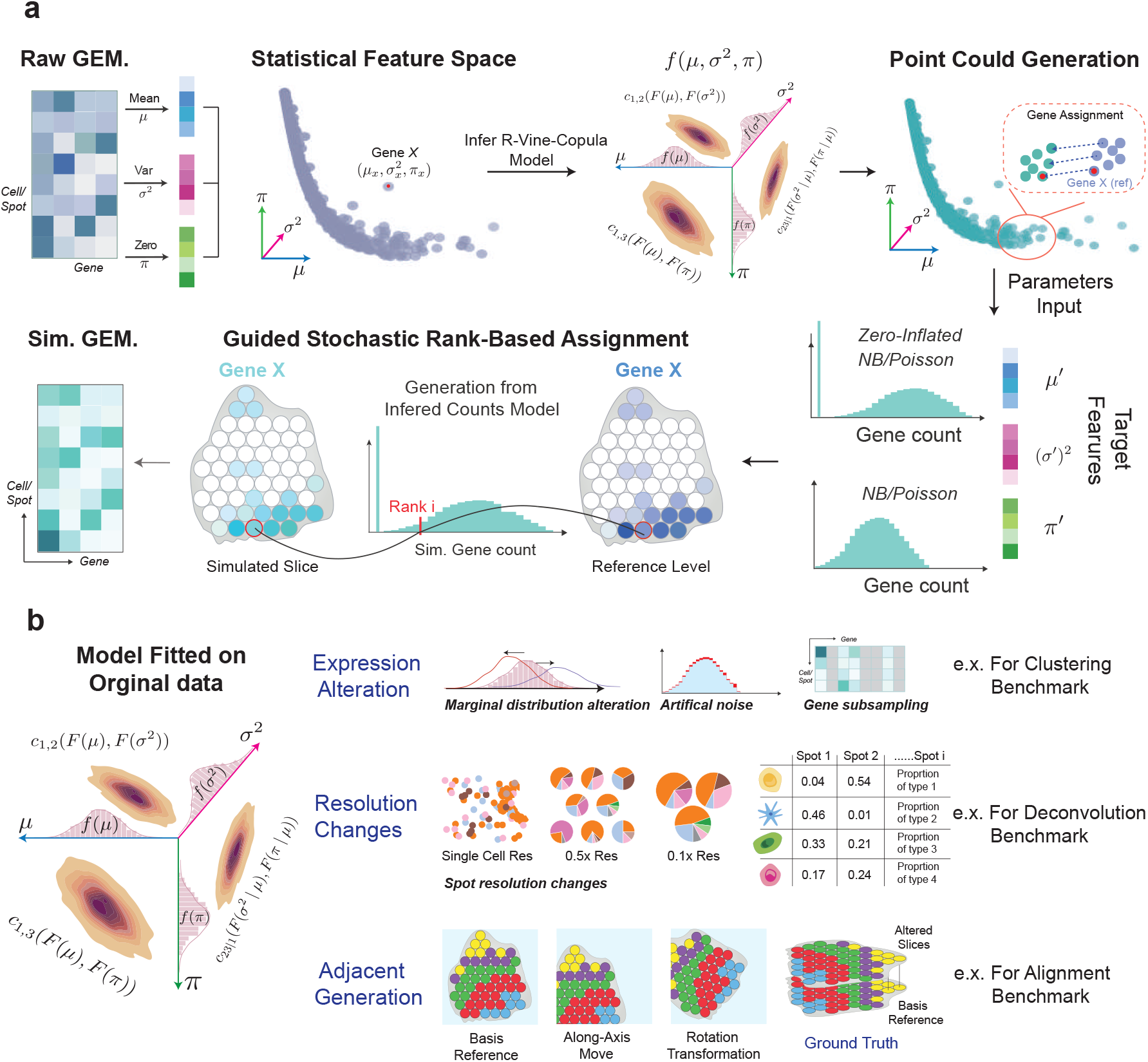
Overview of the FEAST framework. **a**, Core single-slice simulation pipeline. FEAST learns a parameter cloud describing gene-wise mean, variance, and sparsity from a raw gene expression matrix (GEM). A C-vine copula captures the dependency structure among these parameters, enabling the generation of new parameter sets that preserve their joint relationships. These simulated parameters are assigned to genes under a chosen count model to produce synthetic expression profiles, followed by a rank-based mapping to construct the simulated GEM. **b**, Modules for generating benchmark datasets through controlled alteration of expression distributions, spatial resolutions, or spatial transformations to create adjacent slices.

#### Concepts of Parameter cloud and feature space

We first summarize the observed expression data into a compact **parameter cloud** that captures global gene-level behavior. The three-dimensional rectangular coordinate system containing these points of the parameter cloud is called **feature space**. Given the gene expression count matrix 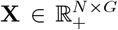, where *N* denotes the number of cells (or spots) and *G* denotes the number of genes, we compute for each gene *g*:

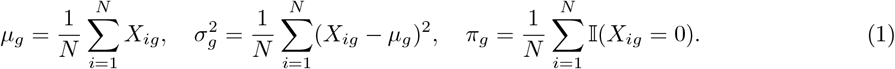

Here, *µ*_*g*_ and 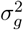 represent the mean and variance of gene *g* across all spots, and *π*_*g*_ denotes the proportion of zero counts (dropout rate). The resulting cloud 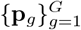 forms an empirical joint distribution over (*µ, σ*^2^, *π*), summarizing how genes vary in expression level, variability, and sparsity within a single slice. This parameter cloud serves as the foundation for our generative model.

#### Marginal models

The next step models each component of the parameter cloud separately to capture the heterogeneity of gene-level statistics across the transcriptome. Each dimension (mean, variance, and sparsity) exhibits distinct statistical behaviors: expression means and variances often show heavy-tailed distributions, while sparsity proportions are bounded within [0, 1] and can be highly skewed. To flexibly represent these diverse patterns, we use mixture models to model the marginal distributions 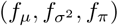, which describe the individual variability of each parameter prior to dependency modeling. Detailed formulations are provided in Extended Methods S1.1.

#### Joint dependency via C-vine copula

To model nonlinear and asymmetric dependencies among (*µ, σ*^2^, *π*), we employ a C-vine copula structure. Copulas allow us to model complex correlations among variables while preserving their fitted marginals. Let 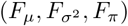 denote the marginal cumulative distribution functions, and define the corresponding uniform variables 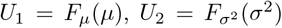, and *U*_3_ = *F*_*π*_(*π*). Under the C-vine construction, the joint density factorizes as (See Fig. 1a):

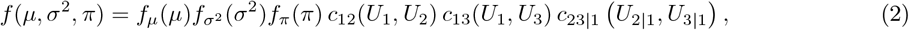

where *c*_*ij*_ denotes the bivariate copula density between variables *i* and *j*, and *U*_2|1_ = ∂*C*_12_(*U*_1_, *U*_2_)*/*∂*U*_1_, *U*_3|1_ = ∂*C*_13_(*U*_1_, *U*_3_)*/*∂*U*_1_ are the conditional uniform variables. Each detailed pair-copula is selected based on the Bayesian Information Criterion (BIC) to balance model fit and parsimony. See details in Extended Methods S1.2.

This copula layer connects the three marginal models into a unified joint distribution *f* (*µ, σ*^2^, *π*) that preserves both the individual variability of each parameter and their interdependence. The resulting model thus represents a generative parameter space for one ST slice, from which new parameter triplets 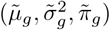 can be sampled. These sampled parameters are subsequently used to simulate gene-level expression profiles that mimic the empirical characteristics of the original data.

#### Synthetic parameters and assignment

From the fitted joint distribution obtained via the C-vine copula, we draw *αG* synthetic parameter triplets 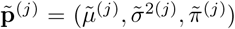, where *α >* 1 controls the number of generated samples relative to the number of real genes *G*. To ensure biologically realistic correspondence, each synthetic triplet is assigned to a real gene through minimum-cost bipartite matching based on standardized features. The optimal one-to-one mapping *π*^∗^ is obtained by solving the Hungarian algorithm. Further methodological details are provided in Extended Methods S1.3.

#### Counts generation and assignment

Having obtained the synthetic parameter triplets 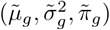 for each gene after assignment via Hungarian algorithm (see Details in Extended Methods S1.3), FEAST then generates spatially distributed expression counts that reflect the gene-specific variability. For each gene, an appropriate probabilistic model is selected from the Poisson, Negative Binomial (NB), Zero-Inflated Poisson (ZIP), or Zero-Inflated Negative Binomial (ZINB) families^21, 28^ based on moments method and data-driven heuristics (See details in Extended Methods S1.4).

For Poisson and Negative Binommial distributions, the estimated parameters will be directly used into model. For ZIP(*λ, π*) and ZINB(*µ, α, π*) distributions, the parameters are estimated by numerically solving a system of equations that align the theoretical moments with the target moments derived from 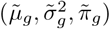.

This moment-matching procedure ensures consistency between the simulated and empirical distributions (see Extended Methods S1.4 for implementation details). The theoretical moments are given by:

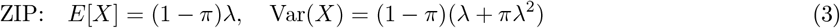

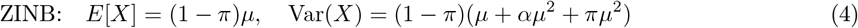

Once each gene’s count distribution is parameterized, FEAST generates synthetic expression counts by random sampling from the selected model and spatially places them via rank-based assignment that preserves local spatial ranks and co-expression module structure (See details in Extended Methods S1.5).

### 2.2 Gene expression and spatial alteration

Let 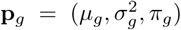 and 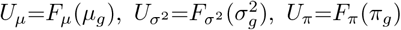. Given altered marginal distributions 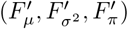, a copula-preserving transformation can be established as follows:

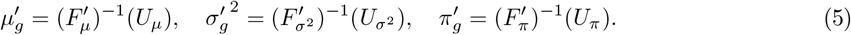

Detailed procedures for expression alteration are provided in Extended Methods S2. For spatial alteration, such as resolution adjustments and positional changes relevant to deconvolution and alignment tasks, please refer to Extended Methods S3-S4 and Fig. 1b.

### 2.3 3D slice interpolation

To reconstruct continuous 3D tissue structures from discrete ST slices, FEAST performs interpolation in the structured feature space rather than directly on noisy expression matrices. This strategy ensures that the interpolated slices preserve realistic gene-level statistical properties, including mean, variance, and sparsity, while maintaining smooth transitions of cellular and molecular patterns across the tissue depth.

The interpolation procedure combines optimal-transport (OT)-based parameter-cloud interpolation with transport-guided spatial coordinates and reference-guided expression reconstruction on the interpolated geometry. By integrating these three components, FEAST generates intermediate slices that are biologically coherent and spatially aligned with adjacent measured sections.

#### Interpolation in feature space

For two adjacent slices with parameter clouds 𝒞_*k*_ and 𝒞_*k*+1_, we define empirical measures over the gene-level parameter triplets 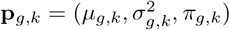.

The parameter cloud 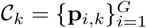 can be viewed as a discrete empirical measure *ν*_*k*_ in ℝ^3^, where each point corresponds to one gene’s statistical signature. For the adjacent slice *k* + 1, the corresponding cloud 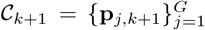 defines another empirical measure *ν*_*k*+1_. Together, these clouds represent the global gene-level landscapes of two consecutive slices, providing a high-level view of molecular variation across tissue depth.

To generate an intermediate slice positioned fractionally between the two, we compute the 2-Wasserstein barycenter of *ν*_*k*_ and *ν*_*k*+1_ for an interpolation coefficient *t* ∈ (0, 1):

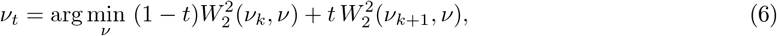

where 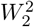 denotes the squared 2-Wasserstein distance between probability measures. Intuitively, *ν*_*t*_ represents the optimal intermediate distribution along the Wasserstein geodesic between slices *k* and *k* + 1, ensuring a smooth and mass-preserving transition between their parameter distributions.

The barycenter *ν*_*t*_ is realized by displacement interpolation under the OT plan **T**^⋆^ with quadratic cost, producing interpolated parameters for each gene index *i* in slice *k*:

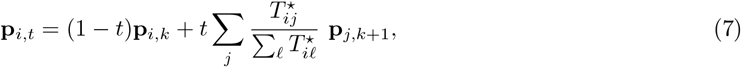

here, **T**^⋆^ is obtained once by minimizing the total quadratic transport cost between the two clouds:

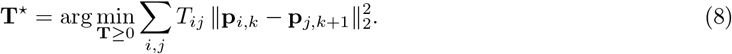

subject to marginal constraints ∑_*j*_ *T*_*ij*_ = *a*_*i*_ and 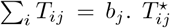 denotes the optimal transport weight between parameter point *i* in slice *k* and point *j* in slice *k* + 1, defining how much “mass” of gene *i* is moved toward gene *j*. The denominator 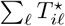 normalizes the mapping for each *i*. In this construction, the index *i* refers to a gene (or feature) in the source slice, *j* indexes its potential matches in the target slice, and *t* controls the interpolation position between the two. The interpolation coefficient *t* does not affect this optimization; instead, *t* determines how far each source point moves along its transport path after **T**^⋆^ is fixed. The resulting **p**_*i,t*_ therefore describes a continuous trajectory between **p**_*i,k*_ and its matched destinations in slice *k*+1.

The resulting interpolated cloud 𝒞_*t*_ = {**p**_*i,t*_}defines the synthetic parameter distribution of an intermediate slice. This parameter cloud serves as the generative foundation for producing virtual ST slices using the single-slice simulation framework, enabling dense 3D continuity for downstream alignment, clustering, and full tissue reconstruction.

#### Alignment-guided coordinates

Following parameter-cloud interpolation, the next step focuses on spatial geometry, interpolating the coordinates of tissue spots between consecutive slices in a way that is consistent with their molecular alignment. We begin with a pre-computed alignment matrix generated by Spateo^16^ 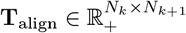, which represents the probabilistic transport plan between spatial spots of adjacent slices *k* and *k*+1.

Each element *T*_align,*ij*_ quantifies the alignment confidence between spot *i* in slice *k* and spot *j* in slice *k*+1. From this matrix, we extract a high-confidence set of correspondences to guide the interpolation of spatial coordinates. This ensures the interpolated geometry preserves biologically meaningful correspondences implied by the molecular transport.

For each high-confidence correspondence, we then generate interpolated coordinates using linear displacement interpolation between the two matched spots:

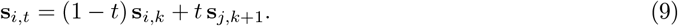

Here, 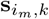 and 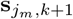 are the 2D spatial coordinates of matched spots in slices *k* and *k*+1, respectively, while *t* ∈ [0, 1] determines the relative position of the interpolated point along the displacement path between them. When *t* = 0, the coordinates correspond exactly to slice *k*; when *t* = 1, they match slice *k*+1; and for 0 *< t <* 1, they yield intermediate spatial positions that ensure smooth geometric continuity across slices.

This interpolation process can be viewed as a spatial analog to the parameter-cloud interpolation: rather than transporting gene-level parameters, it smoothly transports the spatial coordinates of corresponding tissue spots.

#### Ordered query slice

After the gene-level parameters are interpolated through Eq. (7) and the single-slice simulation framework generates corresponding gene count matrices, FEAST can assign these counts onto the interpolated geometry using the previously mentioned rank-based spatial assignment procedure. Because this placement requires reference information from both adjacent slices *k* and *k*+1, FEAST first constructs a unified query slice that aggregates transcriptomic and spatial features from both slices. This query slice serves as a reference layer, encoding expected gene expression patterns across the interpolated coordinates and ensuring that spatial transitions between slices remain smooth and biologically coherent.

To build the gene expression profile for this query slice, FEAST initializes expression at each interpolated spot by blending the aligned source profiles in normalized or log-transformed space. Let *w* = *T*_align, *ij*_ denote the alignment weight between spot *i* in slice *k* and spot *j* in slice *k*+1. The weighted blending is defined as:

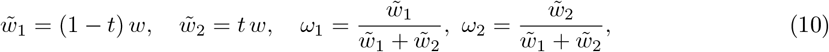

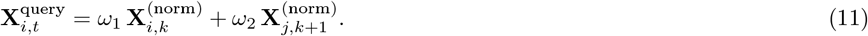

Here, 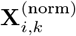 and 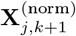 denote the normalized gene expression vectors of matched spots from slices *k* and *k*+1. The coefficients (*ω*_1_, *ω*_2_) define a convex interpolation between these two expression profiles according to the interpolation depth *t*. Because the alignment weight *w* cancels in Eq. (10), the resulting blend depends solely on *t*, producing a smooth transition of expression intensity across depth.

#### Finalization

FEAST then uses this query slice with 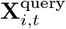 as the spatial-molecular reference for finalizing rank-based gene count assignment, ensuring that each simulated count is positioned according to its anatomically coherent location and biologically consistent expression context.

## 3 Results

### 3.1 FEAST generates high-fidelity spatial transcriptomics slices

To comprehensively assess the performance of FEAST, we benchmarked it against several state-of-the-art spatial and single-cell simulators, including SRTsim, scCube, and the Splatter series. We generated synthetic datasets spanning a broad spectrum of ST platforms and biological contexts, including 10x Visium,29 MERFISH,^30^ OpenST,^31^ Slide-seqV2, Xenium, and Stereo-seq, and systematically evaluated the fidelity of FEAST outputs in capturing gene-level expression statistics, spot-level properties, and spatial expression patterns (Fig. 1; See Extended Methods S5.1).

To isolate the contribution of FEAST’s parameter cloud independent of its generative process, we implemented a streamlined mode that directly employs the extracted parameters without resampling, referred to as FEAST (Raw). The full generative framework is denoted as FEAST (Generative). Benchmark analyses revealed that two modes achieve superior or highly competitive performance across all evaluation criteria (Fig. 2). In preserving fundamental gene-wise statistics, FEAST achieved near-perfect correlation for both gene means and variance estimates. Furthermore, it accurately modeled the characteristic sparsity of ST data, as reflected by minimal Kolmogorov-Smirnov (KS) distances in gene-level zero proportions and spot-level zero proportions. Beyond expression fidelity, FEAST effectively recapitulated complex spatial architectures, achieving Moran’s I and Geary’s C correlations nearly identical to those of the ground-truth reference datasets, thereby demonstrating its ability to preserve both global and local spatial dependencies intrinsic to tissue organization. Our visualization of marker gene expression between experimental and FEAST-generated slices across multiple ST platforms revealed strong concordance in spatial localization and expression intensity patterns (Supplementary Fig. 1). It is worth noting that while SRTsim achieved overall performance comparable to FEAST, this outcome is largely expected because, in its reference-based simulation mode, SRTsim directly derives gene-wise and spatial distribution parameters from the real reference data. In contrast, FEAST re-learns these parameters de novo through its parameter cloud representation. Consequently, FEAST attains similar or superior fidelity while offering substantially greater flexibility and generalizability across spatial platforms and tissue architectures.

**Fig. 2.**
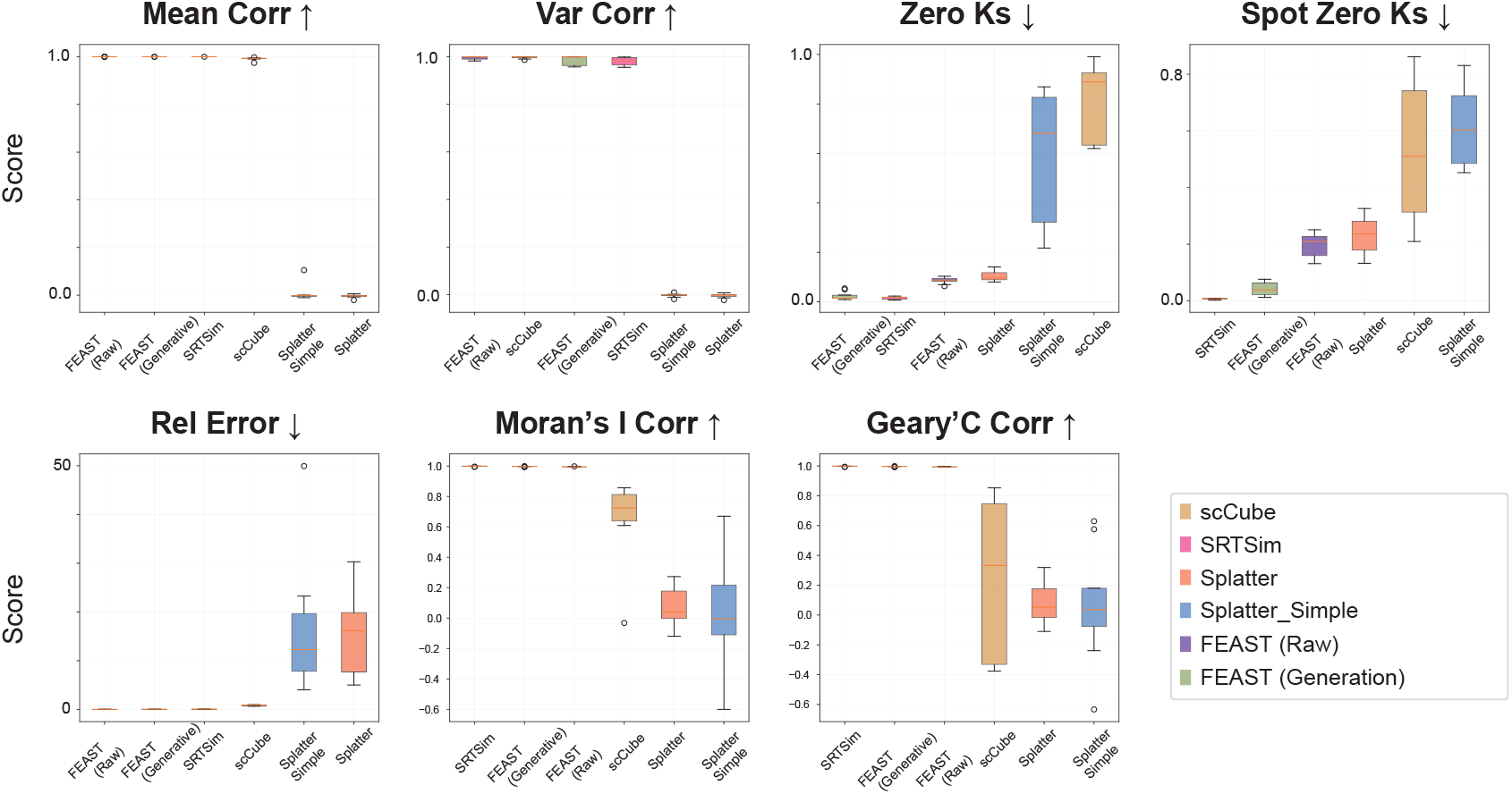
Quantitative benchmarking of FEAST against existing simulators. Boxplots summarize performance across multiple datasets using seven key evaluation metrics: gene-wise mean correlation, variance correlation, zero-proportion Kolmogorov-Smirnov (KS) distance, spot-level zero KS distance, relative count error, and spatial autocorrelation (Moran’s I and Geary’s C). Arrows indicate the direction of improvement for each metric (↑ = higher is better, ↓ = lower is better).

### 3.2 Benchmarking downstream single-slice analysis tools

A central motivation for developing simulation frameworks is to enable systematic and interpretable benchmarking of downstream spatial analysis algorithms. Leveraging FEAST’s controllable data alteration capabilities (Fig. 1b), we generated benchmark datasets with tunable ground-truth perturbations to evaluate clustering and deconvolution performance on single-slice ST data.

#### Robustness analysis of spatial clustering methods

We assessed the robustness of two state-of-the-art spatial clustering algorithms, STAGATE^8^ and GraphST,^10^ by systematically perturbing specific statistical properties of a simulated dataset with known ground truth. Using a 10x Visium human DLPFC slice as the reference, FEAST synthesized altered datasets by independently modifying gene-wise mean expression and sparsity levels while maintaining the underlying joint dependency structure across genes (see Extended Methods S2). The alteration factor of 1.0 corresponds to the unaltered (reference) condition, whereas values below or above 1.0 represent controlled perturbations that are expected to reduce performance as deviation from the original distribution increases. Across the mean-alteration series, STAGATE exhibited higher clustering accuracy (ARI and NMI) and stronger spatial continuity (PAS and CHAOS), maintaining stable performance across a wide dynamic range of mean-expression perturbations. GraphST, in contrast, showed faster degradation as the alteration factor deviated from 1.0. Under sparsity perturbations, STAGATE again demonstrated superior robustness, preserving coherent spatial domains even under high-dropout conditions (Fig. 3). Collectively, these controlled perturbation experiments underscore FEAST’s utility as a principled benchmarking framework: by selectively altering key statistical factors - mean, variance, and sparsity - while preserving spatial organization, FEAST enables mechanistic dissection of model robustness and delineates the operational boundaries of current spatial clustering algorithms.

**Fig. 3.**
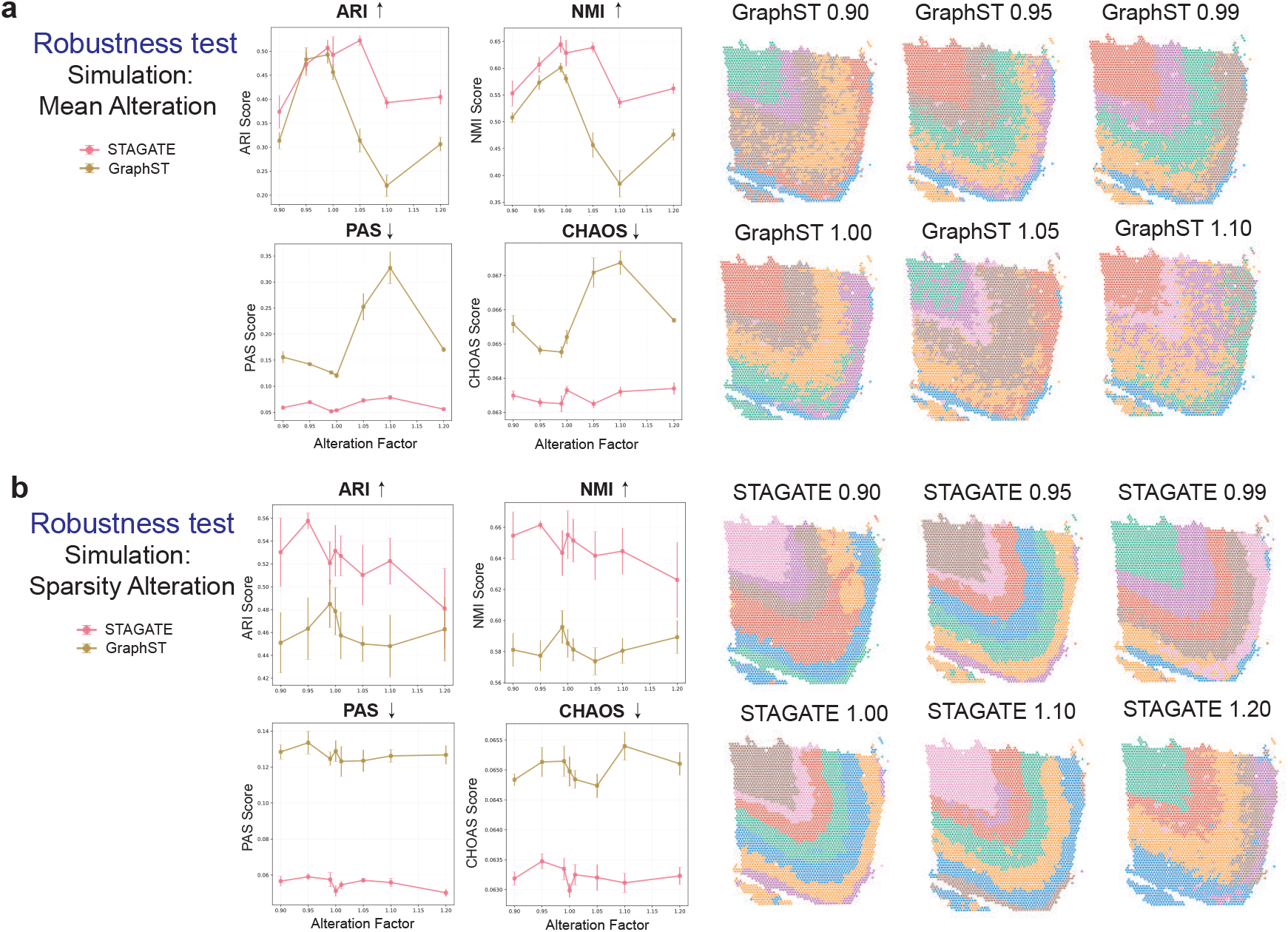
Benchmarking spatial clustering algorithms using controlled expression and sparsity perturbations. **a**, Robustness testing under mean alteration. FEAST generated simulated datasets from a 10x Visium DLPFC slice by systematically modifying gene-wise mean expression, where alteration factors below or above 1.0 represent down-or up-scaling. Line plots show clustering performance of STAGATE (pink) and GraphST (gold) evaluated by Adjusted Rand Index (ARI), Normalized Mutual Information (NMI), and spatial continuity metrics (PAS, CHAOS). Spatial maps visualize clustering outcomes under increasing perturbation magnitudes. The alteration factor of 1.0 (denoted by *) represents the unaltered reference condition. **b**, Robustness testing under sparsity alteration. FEAST introduced controlled changes in dropout rate to assess model stability.

#### Generating ground-truth data for cell-type deconvolution

Robust benchmarking of spatial deconvolution methods requires datasets in which the true cell-type composition of each spot is known, a resource that is rarely available for real ST experiments and is typically approximated using non-spatial pseudo-bulk mixtures. FEAST fills this gap by deriving realistic multi-resolution spot-based datasets from single-cell-resolution spatial maps while retaining full knowledge of the underlying cellular mixtures. Starting from a high-resolution MERFISH slice,^30^ we use FEAST to aggregate single cells into larger synthetic spots on user-defined grids to generate corresponding ST slices at coarser resolutions, recording the exact cell-type proportions per spot as ground truth (Supplementary Fig. 2). In addition, FEAST can introduce controlled gene-level perturbations (e.g., mean and sparsity alterations as in our clustering benchmarks), enabling systematic evaluation of the robustness of current spatial deconvolution tools, such as Cell2location,^32^ across both resolution changes and expression regimes.

### 3.3 Controlled benchmarks for spatial alignment algorithms

Integrating serial tissue sections requires accurate alignment across slices, yet quantitative benchmarking of alignment algorithms remains limited due to the lack of datasets with known spatial correspondences. Most existing evaluations rely on heuristic visual inspection or region-level concordance rather than true spot-to-spot accuracy. FEAST overcomes this limitation by generating paired datasets in which one slice is a precisely controlled geometric transformation, such as rotation, translation, or elastic deformation, of a reference slice while preserving its molecular and spatial context (Extended Methods S3).

To illustrate, we simulated a series of controlled rotations (1°, 5°, 10°, 30°, 45°, and 60°) on a 10x Visium DLPFC slice and systematically compared two state-of-the-art alignment algorithms, Spateo^16^ and SPACEL.^33^ Across multiple evaluation metrics, including Accuracy, Precision, F1 Score, Gene Expression Correlation, and Adjusted Region Accuracy, Spateo consistently outperformed SPACEL (Fig. 4). Spateo maintained near-perfect alignment accuracy and high gene-level correspondence up to a 30° rotation, whereas SPACEL exhibited a rapid decline in performance under moderate to severe transformations (Supplementary Fig. 3).

**Fig. 4.**
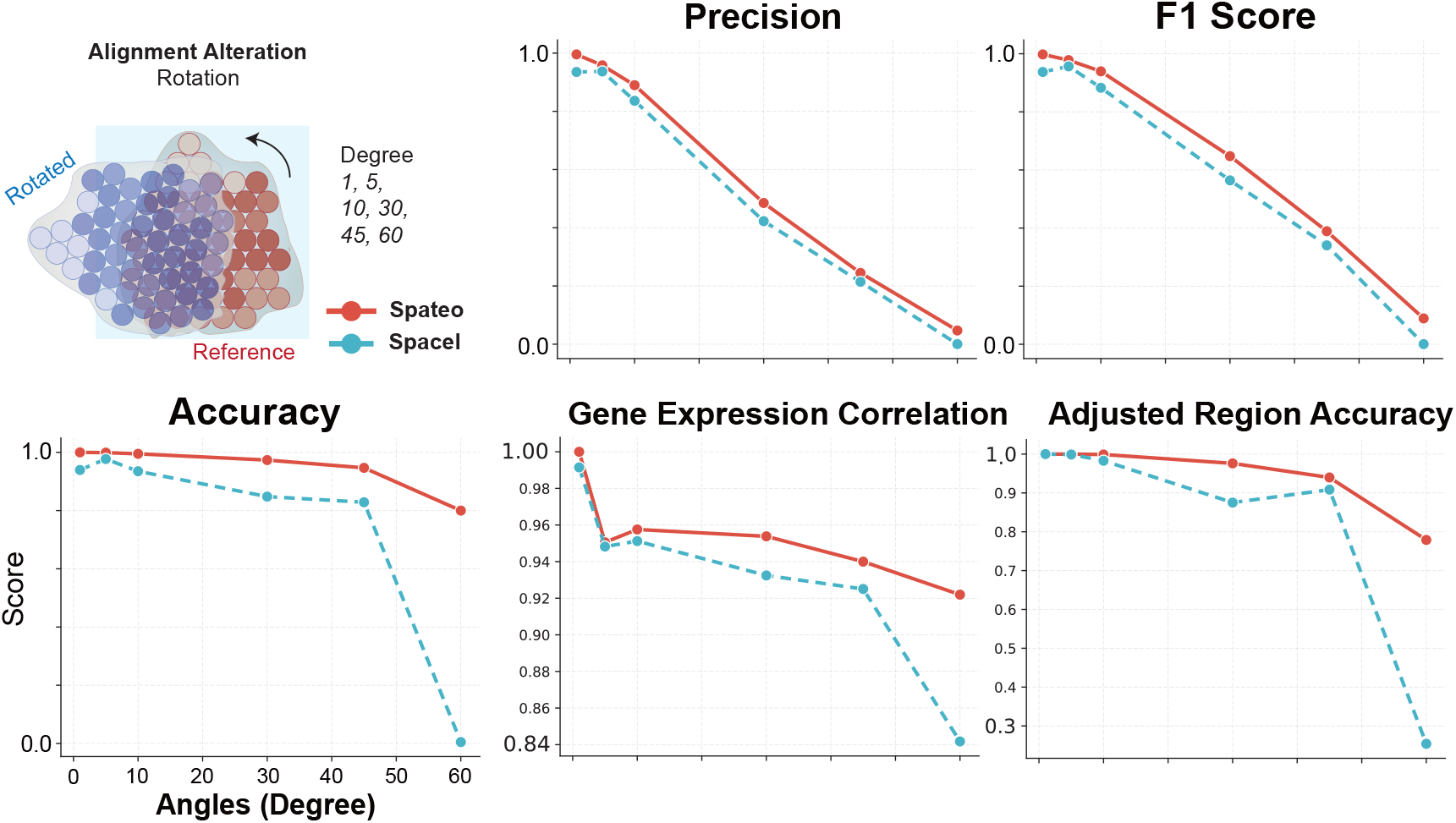
Benchmarking spatial alignment algorithms using controlled geometric transformations. FEAST generated paired slices in which one slice was rotated relative to the reference by defined angles (1°, 5°, 10°, 30°, 45°, 60°) to create datasets with known ground-truth correspondences. Line plots compare the alignment performance of Spateo (red) and SPACEL (blue) across five quantitative metrics: Precision, F1 Score, Accuracy, Gene Expression Correlation, and Adjusted Region Accuracy.

These results demonstrate how FEAST enables quantitative and reproducible assessment of spatial alignment methods under well-defined perturbations, offering a benchmark framework that exposes algorithmic robustness, sensitivity to geometric distortion, and performance boundaries across diverse spatial contexts.

### 3.4 3D interpolation recovers missing slices with high fidelity

Reconstructing continuous three-dimensional tissue volumes from sparsely profiled ST slices remains a key challenge in spatial biology, as most experimental platforms capture only a limited number of serial sections. To address this, FEAST provides a quantitative framework for evaluating interpolation performance under realistic conditions using ground-truth experimental data (Fig. 5a). We assessed its 3D interpolation capability on MERFISH datasets^30^ through a leave-one-out validation design, where middle slices were sequentially withheld and reconstructed solely from their neighboring experimental slices. This setup enables direct and quantitative comparison between interpolated and experimentally measured slices.

**Fig. 5.**
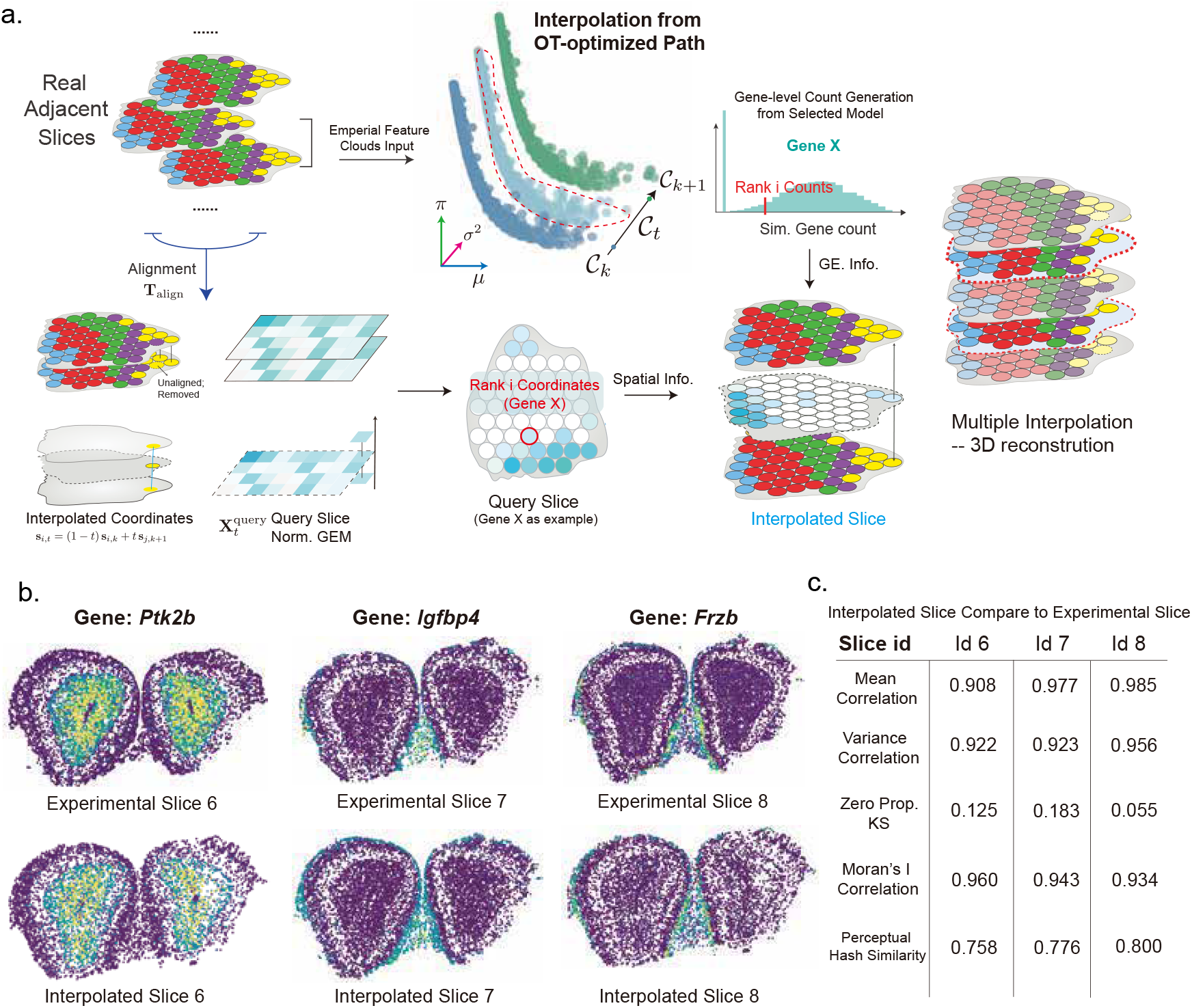
Overview of the FEAST interpolation framework and benchmarking results. **a**, Overview of the FEAST interpolation framework. Details are provided in the Methods section. **b**, Visualization of reconstructed versus experimental slices for representative genes (Ptk2b, Igfbp4, Frzb) from the MERFISH dataset. Interpolated slices closely recapitulate spatial gene expression patterns observed in experimental data. **c**, Quantitative comparison between interpolated and experimental slices (IDs 6-8), showing high concordance across mean and variance correlation, sparsity consistency (zero proportion KS), spatial autocorrelation (Moran’s I), and perceptual hash similarity.

Using five consecutive MERFISH sections (slice IDs 5-9), FEAST reconstructed the omitted middle slices (IDs 6-8) by first aligning adjacent experimental slices, computing optimal transport-based correspondences between parameter clouds of gene-level statistics, and performing smooth interpolation in both spatial and expression domains to regenerate the missing slices. The reconstructed slices recapitulated key anatomical and molecular features, with marker genes showing highly concordant spatial distributions relative to experimental ground truth (Fig. 5b).

Quantitatively, the interpolated slices exhibited strong agreement with experimental data across multiple criteria: mean gene expression correlation and variance correlation both exceeded 0.9, Moran’s I correlation confirmed preservation of spatial autocorrelation structure, and Kolmogorov-Smirnov tests of zero proportion indicated consistent sparsity distributions (Fig. 5c). Additionally, perceptual hash similarity scores corroborated visual fidelity between reconstructed and real slices, validating both molecular and spatial coherence. We further evaluated our interpolation framework by reconstructing full 3D tissue architectures from sparsely sampled slices. Using two MERFISH spinal cord sections spaced 250µm apart as anchors, FEAST-sim generated nine intermediate slices, producing a coherent volumetric reconstruction with smoothly continuous gene-expression patterns (Supplementary Fig. 4). These results illustrate that our approach robustly preserves continuous gene-expression transitions and structural integrity across reconstructed 3D tissues.

Importantly, FEAST’s interpolation module is platform-agnostic and generalizes to any other spatial transcriptomic modalities. By operating on normalized expression matrices and spatial coordinates within a shared parameter-cloud representation, the framework enables unified evaluation and reconstruction across diverse experimental resolutions. Together, these results demonstrate that FEAST can faithfully recover missing spatial layers, facilitating high-fidelity 3D reconstruction of complex tissues from limited or partially sampled spatial data.

## 4 Discussion and Conclusion

FEAST provides a unified and interpretable modeling framework for simulating, benchmarking, and 3D interpolation. By modeling data within a parameter-cloud manifold that encodes gene-level mean, variance, and sparsity while preserving higher-order dependencies, FEAST bridges statistical realism with spatial structure. This framework supports a wide range of applications, including single-slice simulation, ground-truth generation for clustering and deconvolution, alignment benchmarking under controlled geometric transformations, and 3D interpolation for volumetric reconstruction.

