## Supplemental File for "From features to slice: parameter-cloud modeling of spatial transcriptomics for simulation and 3D interpolatory augmentation"

### Supplementary Materials: Supplementary Figures

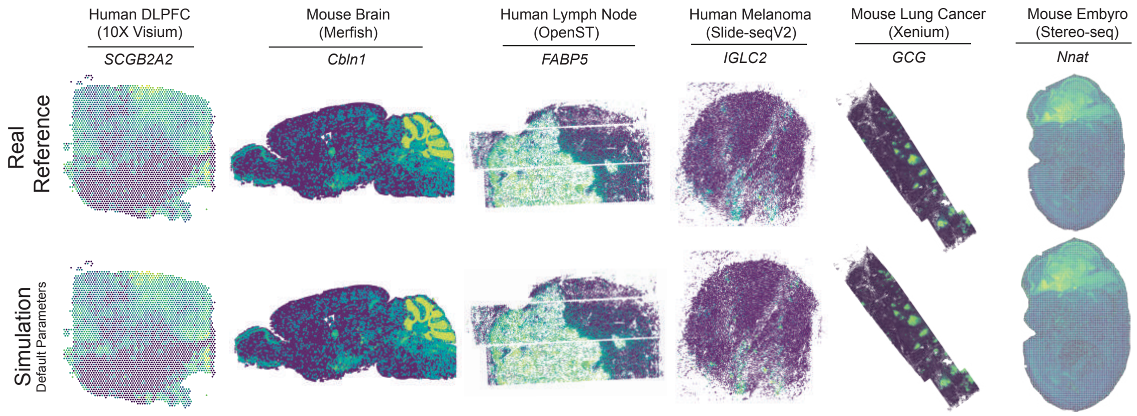


**Supplementary Fig. 1. Visualization of marker gene concordance between real and simulated slices across multiple spatial transcriptomic platforms.** Representative marker genes are shown for six datasets spanning different sequencing technologies: human DLPFC (10x Visium, SCGB2A2), mouse brain (MERFISH, Cbln1), human lymph node (OpenST, FABP5), human melanoma (Slide-seqV2, IGLC2), mouse lung cancer (Xenium, GCG), and mouse embryo (Stereo-seq, Nnat). Simulated slices generated by FEAST under default parameters recapitulate the spatial localization and expression intensity of marker genes observed in real reference datasets, demonstrating high cross-platform fidelity.

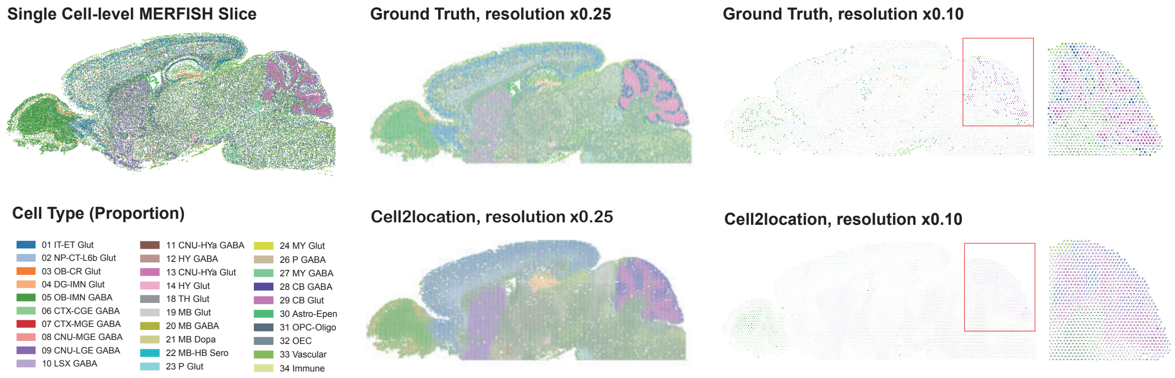


**Supplementary Fig. 2. Benchmarking cell-type deconvolution using simulated datasets across multiple spatial resolutions.** FEAST generates synthetic ST slices with multi-cellular spots derived from single-cell-resolution MERFISH data. it displays simulated ground-truth cell-type proportion maps at two coarser spatial resolutions (0.25× and 0.10× of the original) together with corresponding Cell2location deconvolution results. The simulated slices preserve realistic spatial organization and cell-type mixtures while providing quantitative ground truth for evaluating and calibrating cell-type deconvolution tools across spatial scales.

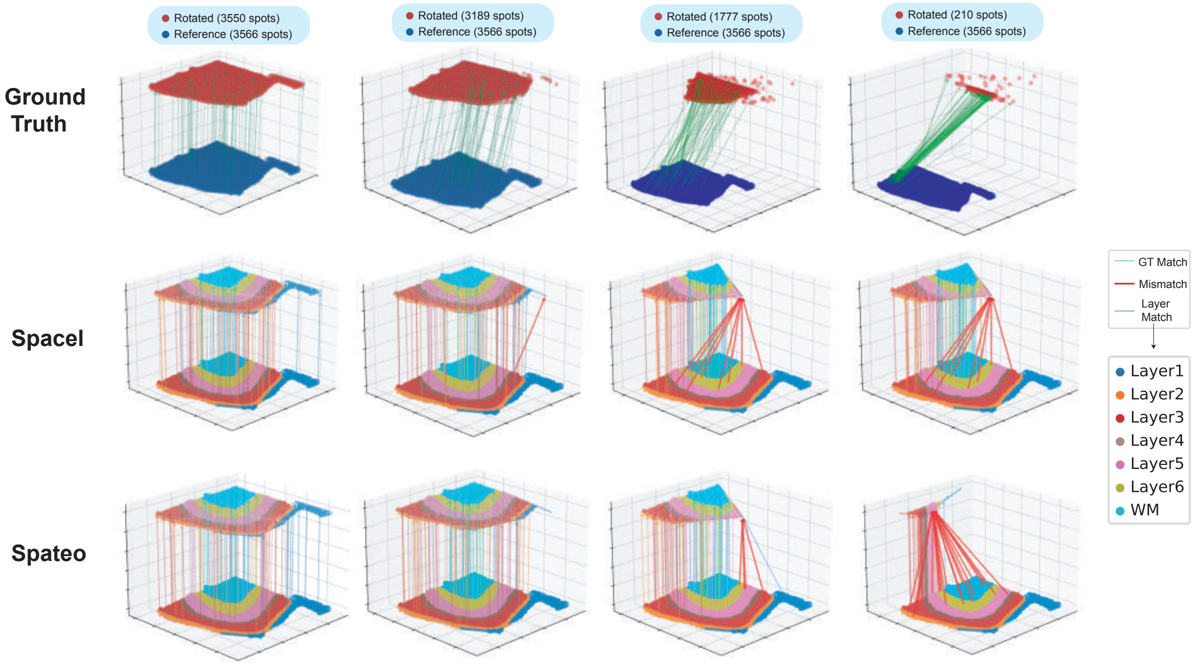


**Supplementary Fig. 3. Illustration of spatial alignment simulation and evaluation.** FEAST generated paired slices with controlled rotational transformations to create ground-truth correspondences for alignment benchmarking. The top row shows ground-truth mappings between reference (blue) and rotated (red) slices, where green lines indicate correct spot-to-spot matches. The middle and bottom rows display alignment results from SPACEL and Spateo, respectively. Correct matches inferred by each algorithm are shown in blue, while incorrect mappings are highlighted in red. Each layer (Layer 1-6 and white matter, WM) is color-coded according to anatomical position. As the rotation angle increases from left to right, mismatches accumulate, particularly in SPACEL, whereas Spateo maintains more accurate cross-layer alignment, demonstrating its robustness to geometric transformations.

### Supplementary Materials: Extended Methods

#### S1 Single-slice Simulation Details

**S1.1 Marginal Model Formulations.** In the marginal mixture models, the mean, variance, and sparsity parameters of each gene are independently modeled to capture their empirical distributions across the dataset. Specifically, the mean and variance parameters are modeled using Student-$t$ mixtures. The density for the mean parameter is shown below; the density for the variance parameter has a similar form and is therefore omitted. The sparsity $\pi \in [0, 1]$ is modeled using Beta mixtures. Each mixture contains up to 10-15 components by default to balance flexibility and computational efficiency:

$$f(\mu) = \sum_{k=1}^{K_\mu} w_k^{(\mu)}\, t(\mu; \nu_k, \theta_k, \sigma_k), \quad t(x; \nu, \theta, \sigma) = \frac{\Gamma\left(\frac{\nu+1}{2}\right)}{\Gamma\left(\frac{\nu}{2}\right)\sqrt{\nu\pi}\sigma}\left(1 + \frac{(x-\theta)^2}{\nu\sigma^2}\right)^{-\frac{\nu+1}{2}}, \tag{12}$$

$$f(\pi) = \sum_{k=1}^{K_\pi} w_k^{(\pi)}\, \mathrm{Beta}(\pi; \alpha_k, \beta_k), \quad \mathrm{Beta}(x; \alpha, \beta) = \frac{x^{\alpha-1}(1-x)^{\beta-1}}{B(\alpha, \beta)}. \tag{13}$$

In the above formulations, $w_k^{(\mu)}$ and $w_k^{(\pi)}$ denote the mixture weights satisfying $\sum_k w_k^{(\cdot)} = 1$. For the Student-$t$ components, $\theta_k$ is the location parameter, $\sigma_k$ the scale, and $\nu_k$ the degrees of freedom controlling tail heaviness. This distribution is well-suited for modeling overdispersed and heavy-tailed behavior commonly observed in gene expression means and variances with decent complexity while remaining accuracy.

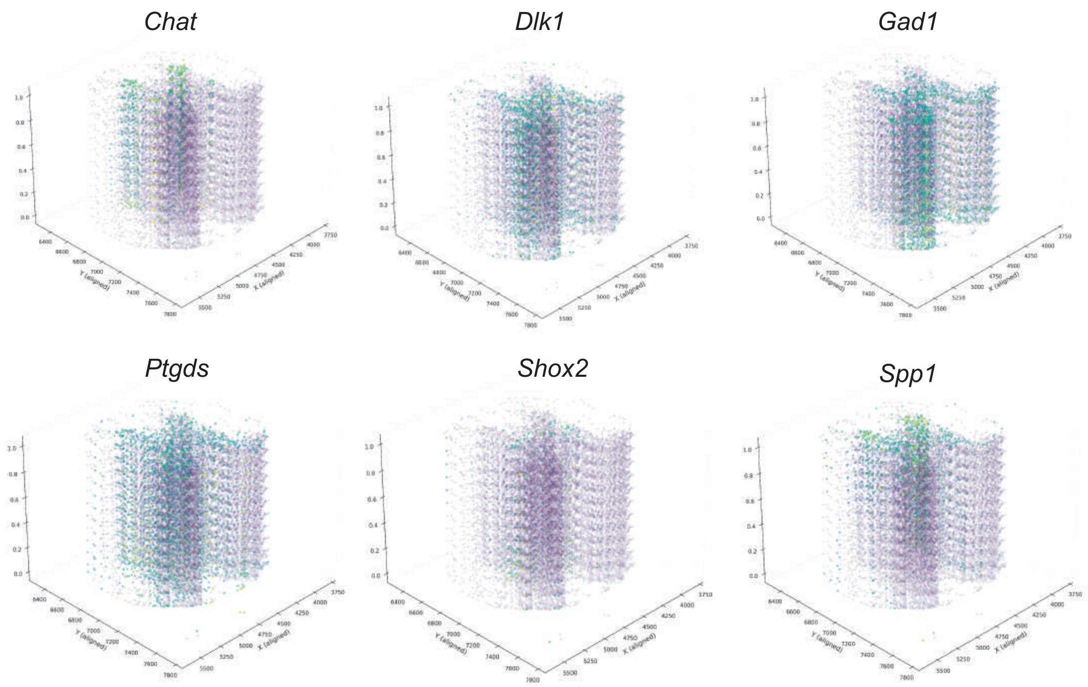


**Supplementary Fig. 4. 3D region reconstruction using two spinal cord MERFISH slices** FEAST generated nine intermediate slices from two reference sections to reconstruct the local 3D spinal cord region. Genes exhibiting similar patterns and expression levels across the two references show smoothly continuous spatial patterns in the interpolated slices. For genes with more variable expression, such as *Spp1*, the model learns and produces a biologically plausible smooth gradient across the reconstructed volume.

For Beta mixtures modeling sparsity: in the Beta mixture, $\alpha_k$ and $\beta_k$ are the shape parameters that determine the skewness and concentration of each component, allowing the model to represent multimodal or asymmetric sparsity patterns across genes. The Beta distribution naturally constrains the sparsity parameter $\pi \in [0, 1]$, making it ideally suited for modeling proportions.

The output of this stage is a set of fitted marginal distributions $(f_\mu, f_{\sigma^2}, f_\pi)$, which describe the individual variability of each parameter independently. However, these parameters are not statistically independent. Genes with high mean expression tend to have higher variance and lower sparsity. To capture such dependencies, a copula-based dependency model is subsequently introduced in the Methods section.

**S1.2 C-Vine Copula Selection Criteria.** Each pair-copula $c_{ij}$ or $c_{ij|k}$ can belong to a different family (e.g., Gaussian, Clayton), allowing flexible modeling of tail dependence and non-Gaussian relationships. The optimal copula family for each pair is selected based on the Bayesian Information Criterion (BIC) to balance model fit and parsimony. This flexibility is crucial because different variable pairs (mean-variance, mean-sparsity, variance-sparsity) may exhibit distinct dependency structures that are best captured by different copula families.

**S1.3 Synthetic Parameter Assignment via Bipartite Matching.** Let $\bar{\mu}_g, \bar{\sigma}_g^2, \bar{\pi}_g$ denote the log-scaled, $z$-scored statistics for gene $g$, and $(\bar{\tilde{\mu}}^{(j)}, \bar{\tilde{\sigma}}^{2(j)}, \bar{\tilde{\pi}}^{(j)})$ denote those for synthetic sample gene $j$. We define the cost matrix as:

$$C_{g,j} = \sqrt{w_\mu(\bar{\mu}_g - \bar{\tilde{\mu}}^{(j)})^2 + w_{\sigma^2}(\bar{\sigma}_g^2 - \bar{\tilde{\sigma}}^{2(j)})^2 + w_\pi(\bar{\pi}_g - \bar{\tilde{\pi}}^{(j)})^2}, \tag{14}$$

where $(w_\mu, w_{\sigma^2}, w_\pi)$ are the feature weights that emphasize mean expression as the dominant matching criterion. By default, these weights are set to prioritize the mean parameter while allowing variance and sparsity to contribute to the matching distance. This weighting scheme reflects the biological importance of the mean expression level in determining gene behavior.

The optimal one-to-one mapping $\pi^*$ between real and synthetic genes is obtained by solving a global minimum-cost assignment problem using the Hungarian algorithm:

$$\pi^* = \arg\min_{\pi} \sum_{g=1}^{G} C_{g,\pi(g)}. \tag{15}$$

This assignment aligns each gene in the original slice with its most similar synthetic counterpart in the parameter cloud, ensuring that downstream simulations maintain realistic per-gene variability and dropout characteristics. The Hungarian algorithm guarantees a globally optimal matching in polynomial time, making this approach computationally tractable even for datasets with thousands of genes.

**S1.4 Count Generation: Theoretical Moments and Parameter Optimization.** For each gene, an appropriate probabilistic model is selected from the Poisson, Negative Binomial (NB), Zero-Inflated Poisson (ZIP), or Zero-Inflated Negative Binomial (ZINB) families based on data-driven heuristics of zero-inflation and overdispersion (default thresholds: $\pi_g > 0.3$ for zero-inflation and $(\sigma_g^2/\mu_g) > 1.5$ for overdispersion). This adaptive selection ensures that genes with sparse, overdispersed expression are modeled more flexibly, while highly expressed and stable genes are represented with simpler distributions. The selection logic is summarized as follows:

- If $\pi_g \leq 0.3$ and $\sigma_g^2/\mu_g \leq 1.5$: use Poisson (equidispersed, no zero-inflation)
- If $\pi_g \leq 0.3$ and $\sigma_g^2/\mu_g > 1.5$: use Negative Binomial (overdispersed, no zero-inflation)
- If $\pi_g > 0.3$ and $\sigma_g^2/\mu_g \leq 1.5$: use Zero-Inflated Poisson (equidispersed, with zero-inflation)
- If $\pi_g > 0.3$ and $\sigma_g^2/\mu_g > 1.5$: use Zero-Inflated Negative Binomial (overdispersed, with zero-inflation)

For ZIP$(\lambda, \pi)$ and ZINB$(\mu, \alpha, \pi)$, where $\alpha$ is the dispersion parameter. The theoretical moments are:

ZIP:

$$E[X] = (1-\pi)\lambda, \quad \mathrm{Var}(X) = (1-\pi)(\lambda + \pi\lambda^2), \quad P(X=0) = \pi + (1-\pi)e^{-\lambda}. \tag{7}$$

ZINB:

$$E[X] = (1-\pi)\mu, \quad \mathrm{Var}(X) = (1-\pi)(\mu + \alpha\mu^2 + \pi\mu^2), \quad P(X=0) = \pi + (1-\pi)(1+\alpha\mu)^{-1/\alpha}. \tag{8}$$

Parameters for each selected model are then inferred from the target triplet $(\hat{\mu}_g, \hat{\sigma}_g^2, \hat{\pi}_g)$ through moment matching and empirical optimization. Specifically, to estimate parameters, we minimize the log-scale squared error between theoretical and target parameters:

$$\begin{aligned}\min_{\boldsymbol{\theta}} \big( &(\log_{10} \mathrm{E}[X;\boldsymbol{\theta}] - \log_{10} \hat{\mu}_g)^2 + (\log_{10} \mathrm{Var}(X;\boldsymbol{\theta}) - \log_{10} \hat{\sigma}_g^2)^2 \\ &+ (\log_{10} P(X=0;\boldsymbol{\theta}) - \log_{10} \hat{\pi}_g)^2 \big)\end{aligned} \tag{16}$$

The log-scale objective is chosen because expression data spans multiple orders of magnitude, and relative errors are more meaningful than absolute errors in this context. The optimization is performed using scikit-learn's numerical optimization routines.

For the simpler NB and Poisson models, parameters are derived directly from the target statistics without numerical optimization:

$$\text{NB:} \quad \mu = \hat{\mu}_g, \quad \alpha = \max\left(0, \frac{\hat{\sigma}_g^2 - \hat{\mu}_g}{\hat{\mu}_g^2}\right) \tag{17}$$

$$\text{Poisson:} \quad \lambda = \hat{\mu}_g \tag{18}$$

The maximum operation in the NB dispersion parameter ensures numerical stability by preventing negative dispersion values.

**S1.5 Spatial Placement via Rank-based Assignment.** The resulting expression profiles are then spatially placed via a rank-based assignment procedure, which preserves local spatial ranks and co-expression module structure observed in the original slice. For each gene, the synthetic counts are sorted and assigned to spatial locations based on their relative ranks in the original slice's spatial distribution. Specifically, if a gene's highest-expressing spot in the original data is at location $i$, the corresponding high-count value in the synthetic data is also placed at an analogous spatial position in the synthetic slice. This rank-based approach ensures that spatial co-expression modules - groups of genes that are highly expressed in nearby regions - are preserved in the simulation while allowing the specific expression magnitudes to vary according to the fitted count distributions.

#### S2 Gene Expression Alteration Details

**S2.1 Parametric Controls for Expression Modification.** To simulate biological conditions or disease-associated perturbations, FEAST provides a set of parametric controls for modifying gene expression characteristics in a controlled and interpretable way.

Mean fold change ($c_\mu$):

$$\log \mu'_g = \log(\mu_g + \varepsilon) + \log c_\mu, \tag{19}$$

where $\varepsilon$ is a small pseudocount (default: $10^{-6}$) added to stabilize computation for genes with zero mean expression. A fold change of $c_\mu = 2$ doubles the mean expression, while $c_\mu = 0.5$ halves it. This control operates in log space to ensure consistent multiplicative scaling across the full expression range.

Dispersion scale ($c_\phi$): the dispersion parameter $\phi_g$ quantifies gene-specific overdispersion beyond the mean. It can be approximated from observed mean and variance statistics as:

$$\sigma_g^2 \approx \mu_g + \phi_g \mu_g^2, \quad \hat{\phi}_g = \max!\Big(0, \frac{\sigma_g^2 - \mu_g}{\mu_g^2}\Big), \tag{20}$$

The adjusted variance under dispersion scaling is:

$$\sigma_g'^{\,2} = \mu'_g + (c_\phi \hat{\phi}_g) \mu_g'^{\,2}, \tag{21}$$

here, $c_\phi > 1$ increases variability (reflecting greater heterogeneity in expression), while $c_\phi < 1$ suppresses variability. This control is particularly useful for simulating biological contexts in which gene expression becomes more or less noisy.

To model changes in zero-inflation (dropout rate) associated with different biological or disease-state perturbations, FEAST adopts a logit-space parameterization of the gene-specific dropout probability $\pi_g$:

$$\operatorname{logit}(\pi'_g) = \delta_\pi + b_\pi, \operatorname{logit}(\pi_g) - \kappa, \log! \frac{\mu'_g}{\mu_g + \varepsilon}, \tag{22}$$

where:

- $\delta_\pi$ is an additive shift on the logit scale, controlling the baseline sparsity level;
- $b_\pi$ is a multiplicative slope on the original $\operatorname{logit}(\pi_g)$, controlling how the gene-specific sparsity scales with its original value;
- $\kappa$ is a coupling parameter linking changes in sparsity to changes in mean expression; The term $-\kappa, \log(\mu'_g/\mu_g)$ ensures that, when mean expression increases and $\kappa > 0$, sparsity naturally decreases, consistent with typical biological behavior.

This three-parameter model provides flexible control over how dropout patterns shift under altered conditions, including both global shifts and expression-dependent adjustments.

#### S3 Alignment Space Alteration

FEAST generates paired ST datasets with controlled geometric transformations, enabling rigorous benchmarking of spatial alignment and registration algorithms. The framework provides exact ground-truth spot-to-spot (cell-to-cell) correspondences while introducing realistic spatial distortions that mimic experimental variability such as tissue rotation, stretching, or local warping.

**S3.1 Workflow Overview.** The alignment simulation workflow consists of three main stages: (a) Expression simulation: a reference slice is first generated using the core FEAST expression simulation module, producing realistic gene expression and spatial distributions. (b) Geometric transformation: controlled spatial operations, including rotation, affine transformation, elastic deformation, or combined transformations, are applied to create a perturbed counterpart of the reference slice. These transformations are parameterized to preserve overall topology while introducing spatial distortions of adjustable intensity. (c) Ground truth construction: the exact mapping between corresponding spots in the reference and transformed slices is recorded as an alignment matrix, providing explicit ground truth for evaluating spatial registration and correspondence algorithms.

**S3.2 Rotation Transformations.** FEAST provides two distinct rotation schemes tailored to different ST technologies:

***Imaging-based rotation.*** For array- or imaging-based platform with fixed spatial coordinates (e.g., 10x Visium, Slide-seq), rotation is applied directly to the coordinate matrix:

$$\mathbf{R}(\theta) = \begin{bmatrix} \cos\theta & -\sin\theta \\ \sin\theta & \cos\theta \end{bmatrix}, \tag{23}$$

$$\mathbf{s}'_i = \mathbf{R}(\theta)(\mathbf{s}_i + \mathbf{c}) - \mathbf{c}, \tag{24}$$

where $\mathbf{s}_i$ denotes the original coordinates, $\mathbf{c}$ is the rotation center, and $\theta$ is the rotation angle in radians. To preserve realistic tissue geometry, an optional filtering step retains only spots within the original spatial boundaries:

$$\mathcal{S}_{\text{filtered}} = \{\mathbf{s}'_i : x_{\min} \le s'_{i,x} \le x_{\max},\ y_{\min} \le s'_{i,y} \le y_{\max}\}, \tag{25}$$

where $(x_{\min}, x_{\max}, y_{\min}, y_{\max})$ define bounding box of the original tissue section.

***Sequencing-based rotation.*** For sequencing-based platforms with irregular spot distributions, FEAST implements a grid-realignment strategy to simulate the discrete capture-array structure:

1. Grid generation: a hexagonal grid is constructed to approximate the original spot density. The minimal inter-spot distance, $d_{\min}$, is estimated using k-nearest-neighbors algorithm. Then:

$$s = \frac{d_{\min}}{2}, \quad h = \frac{d_{\min}\sqrt{3}}{2}, \tag{26}$$

   where $s$ is horizontal spacing and $h$ is vertical offset. The Grid coordinates are:

$$\mathbf{g}_{r,c} = [x_{\min} + 2cs + (r \bmod 2)s,\ y_{\min} + rh], \quad r \in [0, R),\ c \in [0, C), \tag{27}$$

   with $R = \lceil (y_{\max} - y_{\min})/h \rceil + 1$ rows and $C = \lceil (x_{\max} - x_{\min})/(2s) \rceil + 1$ columns.
2. Rotation and assignment: Each original spot coordinate $\mathbf{s}_i$ is rotated using the same rotation matrix as in the imaging-based method to obtain $\mathbf{s}'_i$. The rotated spot is then assigned to its nearest grid point using k-d tree search:

$$\mathbf{s}''_i = \arg\min_{\mathbf{g} \in \mathcal{G}} \|\mathbf{s}'_i - \mathbf{g}\|_2, \tag{28}$$

   where $\mathcal{G}$ is the set of all grid points.
3. Collision resolution: if multiple spots map to the same grid coordinate, only the first assignment is retained, ensuring that the capture layout remains discrete and non-overlapping.

The alignment ground-truth matrix is directly derived from the original cell/spot indices, ensuring that each rotated spot is mapped back to its corresponding original spot.

**S3.3 Edge Artifact Filtering.** Geometric transformations often introduce edge artifacts where rotated tissue extends beyond original boundaries. FEAST provides optional edge filtering to remove spots near boundaries:

$$\mathcal{S}_{\text{clean}} = \left\{\mathbf{s}'_i : x_{\min} + \delta_x < s'_{i,x} < x_{\max} - \delta_x,\ y_{\min} + \delta_y < s'_{i,y} < y_{\max} - \delta_y\right\}, \tag{29}$$

where $\delta_x = \rho(x_{\max} - x_{\min})$ and $\delta_y = \rho(y_{\max} - y_{\min})$ with margin ratio $\rho$ (default: 0.03 or 3%).

#### S4 Spot-Resolution Simulation Details

From single cells resolution slice $\mathbf{Y}_{\text{sc}}$ with annotations $z_n \in \{1, \ldots, K\}$, FEAST synthesizes multi-resolution mixtures with ground-truth compositions. After the simulation of single slice, we could conduct aggregation.

***Multi-resolution aggregation.*** Given a fine level $(\mathbf{X}^{(h)}, W^{(h)}, \mathbf{s}^{(h)})$ and aggregation matrix $\mathbf{A}$,

$$\mathbf{X}^{(\ell)} = \mathbf{A}\mathbf{X}^{(h)}, \quad L^{(\ell)} = \mathbf{A}L^{(h)}, \quad W_{ik}^{(\ell)} = \frac{\sum_u A_{iu}\, L_u^{(h)}\, W_{uk}^{(h)}}{\sum_u A_{iu}\, L_u^{(h)}}, \tag{30}$$

preserving mass and proportions across resolutions.

#### S5 Benchmark Metrics

To rigorously evaluate the performance of spatial transcriptomics methods, we established a comprehensive benchmarking framework that covers four primary tasks: simulation, clustering, alignment, and interpolation. For each task, a suite of quantitative metrics has been designed to assess various facets of algorithmic performance.

**S5.1 Simulation Benchmark Metrics.** These metrics assess the fidelity of simulated spatial transcriptomics data compared to the original data it was modeled from.

To evaluate Distributional Fidelity:

- Mean and Variance Correlation: the Pearson correlation between the gene-wise mean and variance statistics across all genes in the real ($X_{\text{real}}$) and simulated ($X_{\text{sim}}$) data quantifies the similarity of marginal expression distributions.
- Gene and Spot Zero Proportion KS Tests: the Kolmogorov-Smirnov (KS) test is applied to assess the similarity between the distributions of zero proportions calculated per-gene and per-spot in the real and simulated datasets.

Formally, let $\mu_{\text{real},j}$ and $\mu_{\text{sim},j}$ denote the mean expression of gene $j$ in real and simulated data respectively, and $v_{\text{real},j}, v_{\text{sim},j}$ denote their corresponding variances across spots. The Pearson correlations are then computed as:

$$r_{\text{mean}} = \text{corr}\big( (\mu_{\text{real},j})_{j=1}^{g},\ (\mu_{\text{sim},j})_{j=1}^{g} \big), \quad r_{\text{var}} = \text{corr}\big( (v_{\text{real},j})_{j=1}^{g},\ (v_{\text{sim},j})_{j=1}^{g} \big). \tag{31}$$

For the KS test comparing zero-proportion distributions (either per-gene or per-spot), let $F_1(x)$ and $F_2(x)$ denote the empirical cumulative distribution functions (CDFs) of the two groups, the KS statistic is defined as:

$$D = \sup_x \big|F_1(x) - F_2(x)\big|. \tag{32}$$

- Relative Error of Mean Expression: the average relative error between the mean expression of each gene in the real and simulated data.

$$\text{RE}_{\text{mean}} = \frac{1}{g}\sum_{j=1}^{g} \frac{|\mu_{\text{real},j} - \mu_{\text{sim},j}|}{\mu_{\text{real},j} + \epsilon} \tag{33}$$

  and $\epsilon$ means a small constant to avoid denominator to be zero.
- Spatial Autocorrelation: Moran's I and Geary's C are computed for both real and simulated data to ensure spatial patterns are preserved.

Given a spatial weights matrix $W = (w_{ij})$ with total weight $\mathcal{W} = \sum_i \sum_j w_{ij}$, let $x_i$ be the expression (or a summary) at spot $i$, $\bar{x} = n^{-1}\sum_i x_i$, and standardized values $z_i = x_i - \bar{x}$. Moran's I and Geary's C are

$$I_{\text{Moran}} = \frac{n}{\mathcal{W}} \cdot \frac{\sum_i \sum_j w_{ij}(x_i - \bar{x})(x_j - \bar{x})}{\sum_i (x_i - \bar{x})^2}, \tag{34}$$

$$C_{\text{Geary}} = \frac{(n-1)}{2\mathcal{W}} \cdot \frac{\sum_i \sum_j w_{ij}(x_i - x_j)^2}{\sum_i (x_i - \bar{x})^2}. \tag{35}$$

**S5.2 Clustering Benchmark Metrics.** These metrics assess the quality of spatial clustering results.

***Spatial coherence of clusters.*** These metrics evaluate the spatial organization and continuity of the identified clusters.

- CHAOS (CHoice of Anchor points in the Original Space): this metric measures the spatial disorganization of clusters by calculating the average distance from each spot to its nearest neighbor within the same cluster. Lower values suggest more compact and well-organized clusters. After standardizing the spatial coordinates $S$, it is computed as:

$$\text{CHAOS} = \frac{1}{n} \sum_{k \in K} \sum_{i: c_i = k} \min_{j \neq i, c_j = k} d(s'_i, s'_j) \tag{36}$$

  where $K$ is the set of clusters and $d(\cdot,\cdot)$ is the Euclidean distance.
- PAS (Proportion of Adversarial Spots): this metric quantifies the spatial continuity by identifying the proportion of spots whose local neighborhood is dominated by other clusters. A lower PAS indicates better spatial continuity.

$$\text{PAS} = \frac{1}{n} \sum_{i=1}^{n} \mathbb{I}\left[ \sum_{j \in \mathcal{N}_k(i)} \mathbb{I}[c_j \neq c_i] > \frac{k}{2} \right] \tag{37}$$

  where $\mathcal{N}_k(i)$ is the set of $k$-nearest neighbors of spot $i$ (default $k = 10$).

***Standard clustering evaluation.*** When ground-truth labels are available, we employ widely-used metrics to compare the predicted labels ($\hat{Y}$) with the true labels ($Y$):

- Adjusted Rand Index (ARI)
- Normalized Mutual Information (NMI)

Let $n$ denote the number of spots, and $n_{ij}$ represent the entries of the contingency table between true clusters $i$ and predicted clusters $j$. Define row sums $a_i = \sum_j n_{ij}$ and column sums $b_j = \sum_i n_{ij}$. Using $\binom{u}{2} = u(u-1)/2$, the ARI is computed as:

$$\text{ARI} = \frac{\sum_{i,j} \binom{n_{ij}}{2} - \frac{\sum_i \binom{a_i}{2} \sum_j \binom{b_j}{2}}{\binom{n}{2}}}{\frac{1}{2}\Big( \sum_i \binom{a_i}{2} + \sum_j \binom{b_j}{2} \Big) - \frac{\sum_i \binom{a_i}{2} \sum_j \binom{b_j}{2}}{\binom{n}{2}}}. \tag{38}$$

For NMI, given the mutual information $I(Y;\hat{Y})$ and entropies $H(Y), H(\hat{Y})$, we use symmetric normalization as:

$$\text{NMI} = \frac{I(Y;\hat{Y})}{\sqrt{H(Y)\,H(\hat{Y})}}, \quad I(Y;\hat{Y}) = \sum_{i,j} \frac{n_{ij}}{n} \log \frac{n\, n_{ij}}{a_i b_j}. \tag{39}$$

**S5.3 Alignment Benchmark Metrics.** These metrics evaluate the quality of the spatial alignment between two tissue slices.

***Matching accuracy.*** Given a predicted alignment probability matrix $\Pi \in \mathbb{R}^{n_1 \times n_2}$ and a ground-truth matching matrix $G \in \{0,1\}^{n_1 \times n_2}$, where $n_1$ and $n_2$ are the number of spots in each slice, we first derive a binary prediction matrix $P$ by selecting the most likely match for each spot in the first slice:

$$P_{ij} = \begin{cases} 1 & \text{if } j = \arg\max_k \Pi_{ik} \\ 0 & \text{otherwise} \end{cases} \tag{40}$$

From this, we compute the following based on the confusion matrix:

- True Positives (TP): $\sum_{i,j} P_{ij} \cdot G_{ij}$

– False Positives (FP): $\sum_{i,j} P_{ij} \cdot (1 - G_{ij})$
– False Negatives (FN): $\sum_{i,j} (1 - P_{ij}) \cdot G_{ij}$

The performance is then quantified using:

$$\text{Precision} = \frac{TP}{TP + FP} \tag{41}$$

$$\text{Recall} = \frac{TP}{TP + FN} \tag{42}$$

$$\text{F1 Score} = \frac{2 \cdot \text{Precision} \cdot \text{Recall}}{\text{Precision} + \text{Recall}} \tag{43}$$

***Alignment consistency.*** This metric assesses the bidirectionality of the alignment. A consistent alignment should have the forward mapping from slice 1 to slice 2 correspond with the backward mapping from slice 2 to slice 1.

The forward and backward mappings are defined as:

$$f(i) = \arg\max_j \mathit{\Pi}_{ij} \quad \text{(forward: slice 1} \to \text{slice 2)} \tag{44}$$

$$b(j) = \arg\max_i \mathit{\Pi}_{ij} \quad \text{(backward: slice 2} \to \text{slice 1)} \tag{45}$$

Consistency is the fraction of spots in slice 1 that are mapped back to themselves:

$$\text{Consistency} = \frac{1}{n_1} \sum_{i=1}^{n_1} \mathbb{I}[b(f(i)) = i] \tag{46}$$

where $\mathbb{I}[\cdot]$ is the indicator function.

***Region and gene expression concordance.*** These metrics evaluate how well the alignment preserves biological structures and expression patterns.

– Region Accuracy: it measures the proportion of correctly aligned spots that map to the same annotated region. Given region labels $R_1$ and $R_2$ for the two slices and a ground-truth mask $M$ indicating which spots have a valid match:

$$\text{Region Accuracy} = \frac{\sum_{i:M_i=1} \mathbb{I}[r_1^{(i)} = r_2^{(f(i))}]}{\sum_i M_i} \tag{47}$$

– Gene Expression Correlation: for successfully aligned spot pairs, this metric computes the average Pearson correlation of their gene expression profiles. Given expression matrices $X_1, X_2 \in \mathbb{R}^{n \times g}$ for $n$ aligned spots and $g$ common genes:

$$\rho_i = \frac{\sum_{k=1}^{g} (X_{1,ik} - \bar{X}_{1,i})(X_{2,f(i),k} - \bar{X}_{2,f(i)})}{\sqrt{\sum_{k=1}^{g} (X_{1,ik} - \bar{X}_{1,i})^2} \sqrt{\sum_{k=1}^{g} (X_{2,f(i),k} - \bar{X}_{2,f(i)})^2}} \tag{48}$$

The overall correlation is the average over all aligned spots: $\frac{1}{n} \sum_{i=1}^{n} \rho_i$.

**S5.4 Interpolation Benchmark Metrics.** These metrics evaluate the quality of interpolated gene expression data by comparing it to a ground-truth slice.

***Statistical similarity.***

– Mean and Variance Correlation: it computes the Pearson correlation between the gene-wise means and variances of the real ($X_{\text{real}}$) and predicted ($X_{\text{pred}}$) data.
– Zero Proportion KS Test: the Kolmogorov-Smirnov (KS) test is used to compare the distributions of the proportion of zero-expression values per gene in the real and predicted data.

The same definitions of the Pearson correlation and KS statistic as described in Section S5.1 are applied here to compare $X_{\text{real}}$ versus $X_{\text{pred}}$, assessing correspondence in gene-wise means and variances as well as zero-proportion distributions.

***Structural and perceptual similarity.***

- Structural Similarity (Moran's I Correlation): this measures how well the spatial structure of gene expression is preserved by calculating the Pearson correlation between the Moran's I values of each gene in the real and predicted data.
- Perceptual Hash Similarity:[34] this metrics is derived from image similarity metrics. For each gene, we generate a binary "hash" of its spatial expression pattern by downsampling to a small grid (e.g., 8x8) and thresholding at the median expression. The similarity is the average Hamming distance between the hashes of the real and predicted data across all genes. This metric could be reliable and robust to compare the overall spatial similarity, even though the two slices are not in exactly same shape.

For structural similarity, we reuse the Pearson correlation (as in S5.1) computed between per-gene Moran's I values from real and predicted data. For perceptual hashes, let $h^{(g)}_{\text{real}}, h^{(g)}_{\text{pred}} \in \{0,1\}^m$ be the $m$-bit hashes for gene $g$ from real and predicted slices. With Hamming distance $d_H$, define the similarity as

$$S_{\text{pHash}} = 1 - \frac{1}{G}\sum_{g=1}^{G} \frac{d_H\left(h^{(g)}_{\text{real}},\ h^{(g)}_{\text{pred}}\right)}{m}. \tag{49}$$

#### S6 Data Availability

Dataset 1 comprises 12 human DLPFC sections from the spatialLIBD resource (http://research.libd.org/spatialLIBD/); DLPFC 151670 was used for simulator benchmarking, 151676 for clustering evaluation, and 151675 for alignment testing. Dataset 2 consists of 3D MERFISH spatial transcriptomics data with 147 annotated coronal sections from the Allen Brain Atlas (https://alleninstitute.github.io/abc_atlas_access/descriptions/Zhuang-ABCA-1.html); sections ID 6 and 7 were used for simulator benchmarking, section ID 7 for deconvolution testing, and sections ID 5-9 for interpolation evaluation. Dataset 3 contains 19 consecutive slices of annotated 3D OpenST sequencing data from human lymph node (https://www.ncbi.nlm.nih.gov/geo/query/acc.cgi?acc=GSE251926); slices ID 5 and 6 were used for simulator benchmarking. Dataset 4 includes five annotated Stereo-seq slices from the MOSTA mouse atlas (https://www.sciencedirect.com/science/article/pii/S0092867422003993); sample E14_5_E2S2 was used for simulator benchmarking. Dataset 5 contains one Slide-seqV2 slice obtained from the Tencent SpatialOmics database (https://gene.ai.tencent.com/SpatialOmics/dataset?datasetID=119). Dataset 6 includes Human Lymph Node Preview Data sequenced with Xenium from 10x Genomics public datasets (https://www.10xgenomics.com/datasets/human-lymph-node-preview-data-xenium-human-multi-tissue-and-cancer-panel-1-standard) and was used for simulator benchmarking. Dataset 7 includes two MERFISH spinal cord slices from https://github.com/shanmeltzer/Spinal_MERFISH.
